# Nucleosome scaffolding by Brd4 tandem bromodomains in acetylation-dependent chromatin compartmentalization

**DOI:** 10.1101/699967

**Authors:** Michael D. Olp, Vaughn Jackson, Brian C. Smith

## Abstract

Bromodomain binding of acetyl-lysine residues is a crucial step in many epigenetic mechanisms governing transcription. Nearly half of human bromodomains exist in tandem with at least one other bromodomain on a single protein. The Bromodomain and ExtraTerminal domain (BET) family of proteins (BrdT, Brd2, Brd3 and Brd4) each encode two bromodomains at their *N*-termini and are important regulators of acetylation-dependent transcription in homeostasis and disease. Previous efforts have focused on identifying protein acetylation sites bound by individual bromodomains. However, the mechanisms through which tandem bromodomains act cooperatively on chromatin are largely unknown. Here, we first used small angle x-ray scattering combined with Rosetta *ab initio* modeling to explore conformational space available to BET tandem bromodomains. For Brd4, the flexible tandem bromodomain linker allows for distances between the two acetyl-lysine binding sites ranging from 15 to 157 Å. Using a bioluminescence resonance energy transfer assay, we show a clear distance dependence for Brd4 tandem bromodomain bivalent binding of multiply acetylated histone H4 peptides. However, isothermal titration calorimetry studies revealed Brd4 binding affinity toward multiply acetylated peptides does not correlate with the potential for bivalent binding. We used sucrose gradient assays to provide direct evidence *in vitro* that Brd4 tandem bromodomains can simultaneously bind and scaffold multiple acetylated nucleosomes. Intriguingly, our bioinformatic analysis of deposited chromatin immunoprecipitation sequencing data indicates that Brd4 colocalizes with subsets of histone acetyl-lysine sites across transcriptionally active chromatin compartments. These findings support our hypothesis that scaffolding of acetylated nucleosomes by Brd4 tandem bromodomains contributes to higher-order chromatin architecture.

---

Histone-DNA interactions act in a multiscale manner and form the basis for epigenetic processes that dictate transcriptional output over time and space without modifying the underlying genetic code. DNA is organized with eukaryotic nuclei by histone octamers that contain two copies each of H2A, H2B, H3 and H4 (1). At the atomic scale, 145-147 base pairs (bp) of DNA are wrapped around histone octamers to form nucleosomes (2) that control accessibility to DNA replication, repair and transcriptional processes (3, 4). On a larger scale, the nucleosome forms the functional unit of chromatin that organizes eukaryotic DNA in three dimensions (3D) within the nucleus (5). The degree of chromatin compaction partitions eukaryotic genomes between transcriptionally active euchromatin and transcriptionally repressed heterochromatin. These higher-order 3D chromatin structures necessarily bring distant genetic loci into physical proximity, governing complex transcriptional networks (6). For example, enhancers are often located hundreds of kilobases from the promoters they control, and enhancer-promotor looping is critical for 3D chromatin organization that allows enhancers and promoters to communicate within the nuclear space (7). However, the specific protein-histone interactions that lead to formation, maintenance, and disassembly of these critical 3D chromatin structures remains largely unknown.

Which chromatin regions interact depends on the histone post-translational modifications (PTMs) at the sites of interaction. The functional consequences of histone PTMs were first appreciated in the context of lysine acetylation, in which histone hyperacetylation is associated with increased DNA accessibility (8) and active transcription (9, 10). While this correlation was originally attributed to decreased electrostatic attraction between DNA and histones upon charge neutralization of lysine residues by acetylation (11), the functional consequences of acetylation are now known to be more complicated. Discovery of proteins that deposit, remove and specifically bind lysine acetylation, as well as other histone PTMs, has led to the histone language hypothesis in which histone PTMs act in combination to trigger downstream functions in a context-specific manner (12). Histone lysine acetylation is bound by bromodomains, evolutionarily conserved ∼110 amino-acid protein modules (13). Recognition of acetylated nucleosomes by the 46 human bromodomain-containing proteins is involved in diverse transcriptional processes across cell-types in both homeostasis and disease (14–17).

Anti-proliferative and anti-inflammatory effects associated with bromodomain inhibition and genetic knockdown have driven recent drug discovery efforts (18, 19). Selective inhibitors of bromodomain and extraterminal domain (BET) family of bromodomains (BrdT, Brd2, Brd3 and Brd4) were discovered in 2010 (20, 21) and since have entered clinical trials as potential treatment for cancer (22–27) and inflammation-driven disease (28, 29). However, the molecular mechanisms underlying the potential therapeutic effects of bromodomain inhibitors are poorly understood. As isolated bromodomains harbor relatively weak affinity toward monoacetylated histone tail peptides *in vitro* (*K*_d_ = 10 *µ*M - 1 mM) (13), multivalency is emerging as an increasingly crucial concept in bromodomain binding of modified chromatin (30, 3 1). One category of potential multivalent interactions is the recognition of multiple sites on one histone tail by an individual bromodomain. For instance, the *N*-terminal bromodomains (BD1) of Brd4 and BrdT cooperatively bind two adjacent acetyl-lysine residues (*i.e.* lysines 5 and 8) on the histone H4 tail (13, 32) with 3-to 20-fold tighter affinity compared to either acetylation in isolation (33, 34) suggesting that multiple neighboring acetylation sites are necessary to recruit these bromodomains to chromatin.

Another potential category of multivalent bromodomain-nucleosome interactions is possible through the encoding of proteins containing multiple (tandem) bromodomains in a single polypeptide, a feature observed in yeast to mammalian genomes including 11 human proteins. Early studies of tandem bromodomain proteins in yeast demonstrated that subunit 4 of the chromatin structure-remodeling complex (Rsc4) and bromodomain-containing factor 1 (Bdf1) each require their tandem bromodomains for full function (35, 36). The first crystallographic study of a tandem bromodomain proposed that the bromodomains of human transcription initiation factor TFIID subunit 1 (Taf1) can bind multiple acetylation sites on the same histone H4 tail (37). Recently, studies of human polybromo-1 (Pbrm1) that contains six bromodomains in tandem have led to a similar hypothesized model in which subsets of the Pbrm1 bromodomains work in concert to bind and position multiply acetylated nucleosomes (38–40). Similarly, Brd4 tandem bromodomains associate with nucleosomes hyperacetylated on histones H3 (lysines 9, 14, 18, 23 and 27) and H4 (lysines 5, 8, 12, 16 and 20) with tighter affinity compared to nucleosomes hyperacetylated at either H3 or H4 in isolation (41). Proteins containing multiple bromodomains have also been implicated in the regulation of higher-order chromatin structure. The yeast and drosophila Brd4 homologs Bdf1 and Female sterile homeotic protein (Fsh) are associated with the maintenance of boundaries separating transcriptionally active and inactive chromatin regions (42, 43). The observation that a Brd4 knockdown leads to widespread unfolding of subnuclear chromatin structure has led to the hypothesis that Brd4 functions to scaffold separate nucleosomes and bring them together within the nuclear space (44).

Members of the BET family each contain two bromodomains in tandem at their *N*-termini (referred *N*-to *C*-terminally as BD1 and BD2, respectively) and represent the most targeted subset of human bromodomains in clinical trials (45). The BET family member Brd4 is especially critical for initiation and elongation by RNA Polymerase II (RNAP II) (46) and enhancer-mediated transcription (47, 48), particularly in the contexts of inflammation (49, 50) and Myc-driven cancer (51, 52). While the specificities of individual recombinant BET bromodomains toward histone lysine acetylation have been explored (13, 32–34), it was unknown if the tandem BET bromodomains are capable of simulataneously binding and scaffolding two acetylated nucleosomes, which would have broad implications on how 3D chromatin organization is established in an acetylation-dependent manner. Here, we directly test different possible mechanisms of multivalent BET tandem bromodomain engagement of nucleosomes using biochemical, structural, biophysical, and bioinformatic techniques. We used small-angle X-ray scattering (SAXS) and Rosetta protein modeling to demonstrate that BET tandem bromodomains are linked by relatively long and disordered amino-acid sequences compared to the tandem bromodomains in the non-BET family member Taf1. This long and disordered linker suggests that BET tandem bromodomains can span longer range chromatin interactions. We used a NanoBRET-based assay in combination with isothermal titration calorimetry (ITC) affinity measurements to determine the distance requirements for the Brd4 tandem bromodomain conformational changes that occur in response to binding multiply acetylated H4 peptides. Our bioinformatic analysis of deposited chromatin immunoprecipitation sequencing (ChIP-seq) data indicates that Brd4 colocalizes with subsets of histone acetyl-lysine sites across annotated topologically associated chromatin domains. We also provide the first direct evidence for physical scaffolding of multiple acetylated nucleosomes by Brd4 tandem bromodomains using *in vitro* sucrose gradient binding assays. These observations allow us to hypothesize a role for Brd4-mediated nucleosome scaffolding in the maintenance of higher-order chromatin structure.

## Results

### BET bromodomains are separated by flexible linkers that afford a high degree of conformational freedom

We hypothesize that the range and relative occupancy of intersite distances between tandem bromodomain acetyl-lysine binding sites dictate the types of multivalent nucleosome interactions accessible to tandem bromodomains. These multivalent interactions can broadly be classified into two possibilities that require very different distances between acetyl-lysine binding sites: 1) within one or between two histone tails in one nucleosome (intranucleosome) or 2) between histone tails on two different nucleosomes (internucleosome) (Figure 1). To determine the sizes and shapes adopted by tandem bromodomains of all four BET proteins and Taf1 (Figure S1) in solution, small angle X-ray scattering (SAXS) studies were performed. Atom pair distribution distance distribution (P(*r*)) functions and Kratky representations were calculated to assess protein size/shape and flexibility, respectively. The P(*r*) function is a histogram of distances between all possible pairs of atoms within a particle (53). For Brd4 (aa 36-460), the P(*r*) function is asymmetric around a single peak indicating elongated cylindrical structures with a maximum diameter (*D*_*max*_) of 201 Å (Figure 2A). The Kratky plot allows for qualitative assessment of protein flexibility. In general, scattering intensity from a solid body decays at high angles (q) resulting in a bell-shaped plot whereas scattering intensity from an extended thin chain presents as a plateau followed by a monotonic increase (53). The Kratky plot of Brd4 demonstrates an initially bell-shaped curve that increases monotonically at higher angles (Figure 2B) indicating a combination of ordered and disordered protein regions, consistent with the presence of folded bromodomains connected by a flexible linker sequence. These elongated structures were conserved across BET bromodomains as SAXS analysis of BrdT, Brd2 and Brd3 displayed similarly shaped P(*r*) and normalized Kratky distributions to those of Brd4 (Figure S2A-F). However, not all human tandem bromodomains adopt similar elongated structures. SAXS measurements of the Taf1 tandem bromodomains showed a relatively symmetrical P(*r*) function around two maxima and a Kratky plot with two peaks in the small-q range (Figure S2G). In addition, the *R*_*g*_ and *D*_*max*_ values calculated for the Taf1 bromodomains (30.5 ± 0.5 and 103 ± 5 Å, respectively) were considerably smaller than those of BET tandem bromodomains (48.6-55.4 and 181-200 Å, respectively). These findings are consistent with a relatively globular and ordered Taf1 tandem bromodomain organization compared to the BET tandem bromodomains. The observed differences likely result from the relatively long inter-bromodomain linkers as defined in UniProt (54) for BrdT (169 aa, UniProt ID: Q58F21), Brd2 (199 aa, P25440), Brd3 (201 aa, Q15059) and Brd4 (219 aa, O60885) compared to that for Taf1 (51 aa, Q8IZX4).

**Fig. 1.**
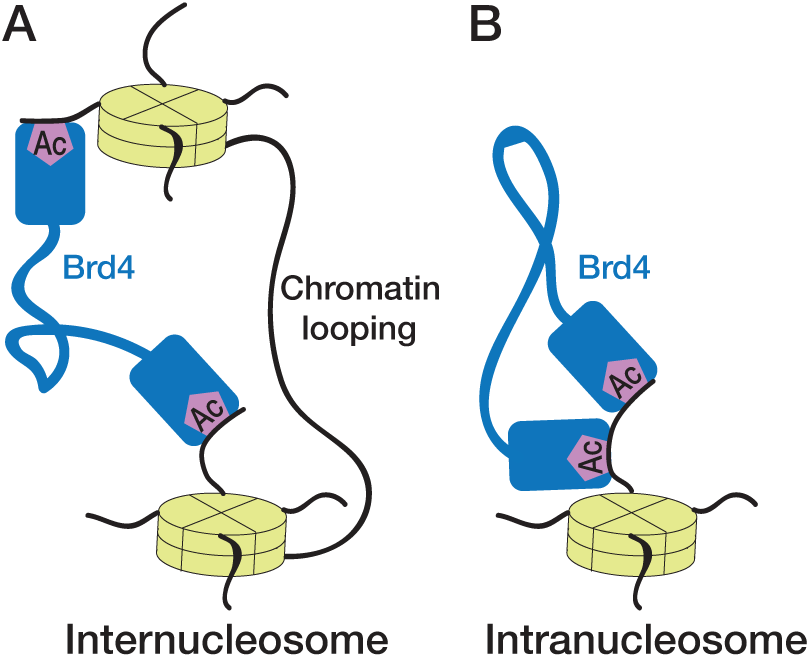
Potential multivalent interactions of tandem bromodomains with chromatin. (A) Intranucleosome interactions (*e.g.* between two acetylation sites on one histone tail) require shorter distances between acetyl-lysine binding sites. (B) Internucleosome interactions (*e.g.* between two distinct promoters) require the longer distances between acetyl-lysine binding sites

**Fig. 2.**
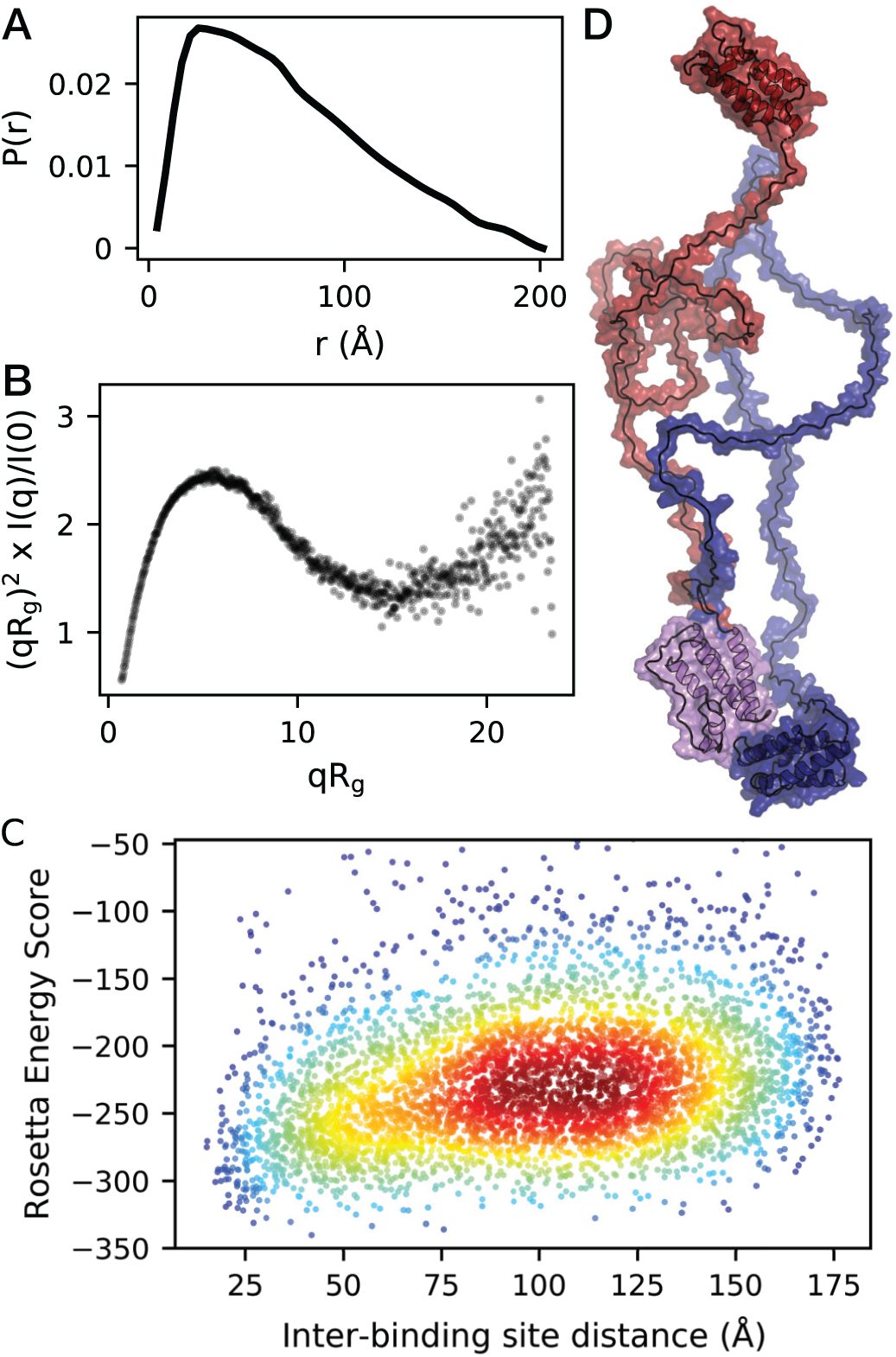
Brd4 bromodomains are separated by flexible linker allowing conformational freedom. (A) P(r) distribution resulting from SAXS measurements of the Brd4 tandem bromodomains (aa 36-460). (B) Rg-normalized Kratky plot of Brd4 tandem bromodomain (aa 36-460) SAXS intensity. (C) Rosetta Energy Score vs. inter-bromodomain binding site distance plot of the top 5,000 models passing (*R*_*g*_ and *D*_*max*_ filters determined by SAXS analysis. Distances were measured using the side chain-NH2 groups of conserved bromodomain acetyl-lysine binding pocket Asn residues 140 (BD1) and 433 (BD2). (D) Representative Rosetta models demonstrating the range of inter-bromodomain acetyl-lysine binding site distances displayed by structures with Rosetta energy scores below the mean of the top 5,000 models (−202.4). Distances between N140 of Brd4-BD1 and N433 of Brd4-BD2 range from 15.2 Å (blue structure) to 176.3 Å (red structure).

When considering potential intranucleosome and internucleosome interactions for tandem bromodomains (Figure 1), the occupancy of distances between the two acetyl-lysine binding sites is particularly important. To computationally model this distance population, *ab initio* modeling of the Brd4 inter-bromodomain linker was performed using the Rosetta Floppy-Tail application. Output models were filtered using the SAXS experimental (*R*_*g*_ = 55.4 ± 5.5 and *D*_*max*_ < 201 Å; Figure S3) values as constraints. Inter-bromodomain acetyl-lysine binding site distances were measured for each filtered output model using the sidechain-NH_2_ groups of the conserved Asn residues (Brd4 N140 and N433) in each bromodomain known to form a critical hydrogen bond with the acetyl-lysine oxygen (13, 55, 56). While a continuous distance distribution ranging from 15 to 157 Å was observed between the two Asn residues, no Rosetta energy convergence within the allowed *R*_*g*_ and *D*_*max*_ ranges was observed (Figure 2C). These results indicate that there is little to no energy barrier in the Brd4 tandem bromodomains accessing this range of inter-binding site distances (Figure 2D).

### Brd4 tandem bromodomains can access internucleosome interactions

Given the predicted long flexible linker between BET bromodomains, we next investigated whether the Brd4 tandem bromodomains can simultaneously bind two separate nucleosomes (Figure 1A). This would be a novel mechanism for scaffolding acetylated chromatin and maintaining 3D chromatin organization through acetylation-dependent chromatin looping. As nucleosomes are ∼57 Å wide, we hypothesize that inter-bromodomain binding site distances >57 Åare necessary for tandem bromodomains to participate in efficient long-range internucleosome interactions. Greater than 86% of the apo structures calculated by our SAXS-based Rosetta modeling display inter-bromodomain acetyl-lysine binding site distances >50 Å (Figure 2C). To measure the inter-bromodomain bindings site distance distribution experimentally, electron paramagnetic resonance (EPR) probe was designed in which a nitroxide spin label (TEMPO) was attached to an inhibitor (JQ1) (20) that binds all BET acetyl-lysine binding sites with nanomolar affinity (‘JQ1-TEMPO’; Figure S4A). Although a BET bromodomain-targeted spin label had not been previously reported, we and others have shown that the location on JQ1 where TEMPO was attached tolerates modification in a manner that does not affect bromodomain binding (33, 34). JQ1-TEMPO binding to the Brd4 tandem bromodomains was shown by decreased amplitude (increased width) of the resonance lines in the continuous wave (cw) EPR spectrum (Figure S4B). Our intention was to collect double electron-electron resonance (DEER) measurements of JQ1-TEMPO simultaneously bound to both Brd4 acetyl-lysine binding sites as a reporter of the native distribution of the inter-bromodomain binding site distances (Figure S4C). However, after titrating JQ1-TEMPO to saturation, as monitored by continuous-wave EPR (Figure S4C), no DEER signal was detected. This result indicates that the nearest-neighbor distances of JQ1-TEMPO, when simultaneously bound to both Brd4 bromodomains, was beyond the instrument detection limit of ∼60 Å. These EPR data are consistent with the vast majority of Brd4 tandem bromodomain binding sites residing >60 Å from each other and therefore the ability of Brd4 tandem bromodomains to access internucleosome interactions.

To directly test the ability of Brd4 tandem bromodomains to simultaneously bind separate nucleosomes, sucrose gradient binding assays were performed with the tandem Brd4 bromodomains and mono-nucleosomes purified from calf thymus and chick erythrocytes (Figure S5). Binding to multiple mononucleosomes should increase their apparent size and therefore their sedimentation rate, shifting them down in a sucrose gradient (Figure S5). Consistent with the ability of Brd4 to access acetylation-dependent internucleosome interactions, addition of Brd4 (aa 36-460) to calf thymus mono-nucleosomes harboring high levels of lysine acetylation (57, 58) resulted in an increased sedimentation rate indicative of larger effective size similar to polynucleosomes relative to control without Brd4 addition (Figure 3A-B).

**Fig. 3.**
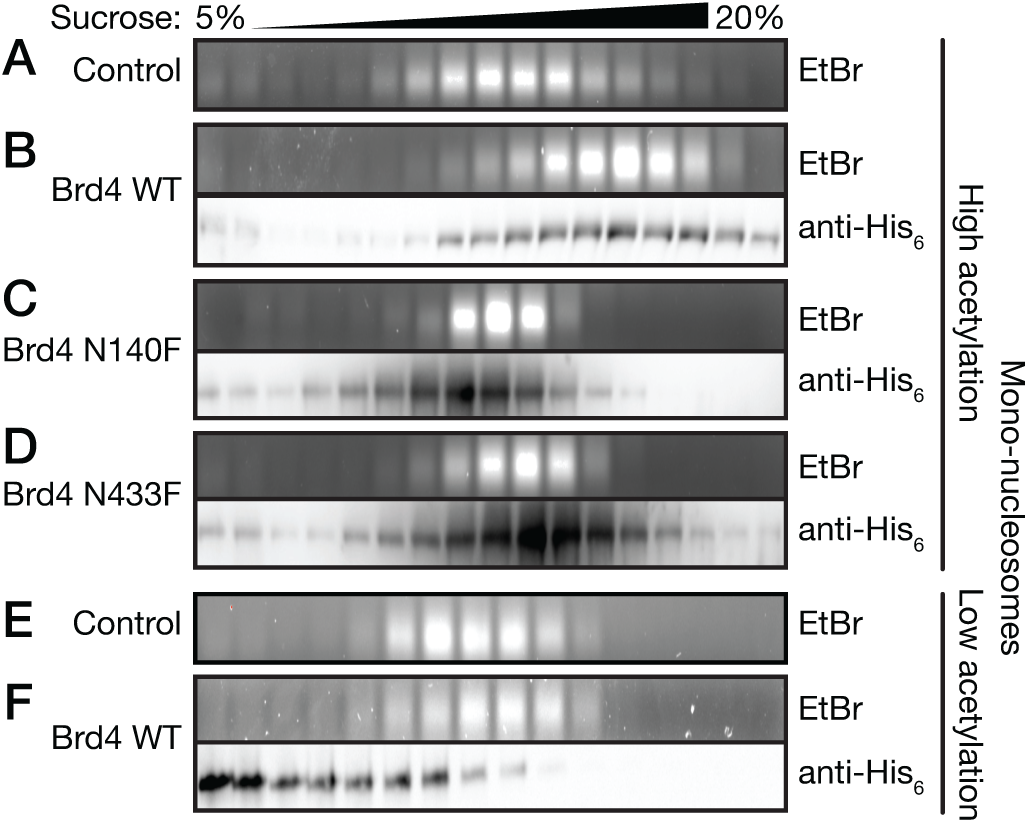
Brd4 tandem bromodomains scaffold acetylated nucleosomes *in vitro*. Sucrose gradient binding assays of the Brd4 tandem bromodomains toward chick erythrocyte (low acetylation) and calf thymus (high acetylation) mono-nucleosomes. Nucleosome-containing sucrose gradient fractions were monitored by agarose gel electrophoresis and ethidium bromide staining of DNA fragments after nucleoprotein digestion. Brd4-containing fractions were detected using immunoblot detection of the *N*-terminal His_6_ tag. (A-B) Wild-type Brd4 tandem bromodomains physically associate with calf thymus mono-nucleosomes (high acetylation) and increase their sedimentation rate relative to control. (C-D) Single bromodomain inactive mutants N140F (BD1) and N433F (BD2) physically associate with acetylated calf thymus mono-nucleosomes but do not increase their sedimentation rate relative to control. (E-F) Wild-type Brd4 tandem bromodomains (aa 36-460) do not increase the sucrose gradient sedimentation rate of chick erythrocyte mono-nucleosomes (low acetylation) relative to controls and show only weak physical association.

We next investigated whether Brd4-mediated nucleosome scaffolding requires acetyl-lysine binding activity within both bromodomains or could instead be explained by DNA-binding activity. Consistent with the acetylation-dependence of Brd4 internucleosome interactions, no increase in nucleosome sedimentation rate was observed when either of the conserved Asn residues required for Brd4-BD1 (N140) or Brd4-BD2 (N433) acetyl-lysine binding (13, 55, 56) were mutated to phenylalanine (59) (Figure 3C-D). Similarly, when Brd4 (aa 36-460) was combined with chick erythrocyte mono-nucleosomes that natively harbor very low acetylation levels (60), no increase in nucleosome sedimentation rate was observed and the majority of the Brd4 protein remained at the top of the sucrose gradient (Figure 3E-F), indicating very weak affinity between Brd4 and either the histones or DNA in chick erythrocyte nucleosomes. Overall, our results demonstrate that the tandem Brd4 bromodomains scaffold nucleosomes *in vitro* in a manner that requires histone acetyl-lysine binding by both bromodomains and that DNA-binding is not sufficient to mediate internucleosome interactions.

### Bivalent binding of multiply acetylated histone H4 tails by Brd4 tandem bromodomains does not result in tighter affinity

Another potential mechanism of multivalent tandem bromodomain interaction with chromatin is bivalent binding of a single histone tail acetylated at multiple sites. This bivalent interaction could strengthen affinity and recruit tandem bromodomains only to chromatin regions hyperacetylated at specific lysine residues. Although our DEER results (Figure S4) indicated that the apo Brd4 tandem bromodomains reside ∼60 Å from each other, our SAXS-guided Rosetta modeling indicated the two Brd4 acetyl-lysine binding sites can access inter-binding site distances as small as 15 Å (Figure 2C), theoretically allowing bivalent interaction with two acetylation sites spaced as closely as 4 amino-acids on an extended histone tail. To investigate the ability of the Brd4 tandem bromodomains to bivalently engage multiply acetylated histone tails, triacetylated histone H4 peptides harboring acetylation at lysine residues 5 and 8 were synthesized with a third acetylation site placed at either lysine 12, 16 or 20. We and others have shown that Brd4-BD1 preferentially binds peptides diacetylated at lysines 5 and 8 (13, 33, 34) and hypothesized that Brd4-BD2 would simultaneously bind to the third acetylation site at lysine 12, 16 or 20 only when the distance between the lysine sites was long enough to accommodate both bromodomains. To test this hypothesis, a previously reported cellular NanoBRET assay was adapted for *in vitro* experiments using recombinant HaloTag-Brd4-BD1_BD2-NanoLuc (aa 44-460) protein. Bivalent engagement of both bromodomains in this Brd4 construct by a single ligand would bring the NanoLuc tag and fluorophore-labeled HaloTag in physical proximity, resulting in increased BRET signal. Addition of either the H4K5/8-diacetyl or H4K5/8/12-triacetyl peptide did not result in increased NanoBRET signal. In contrast, the H4K5/8/16-triacetyl and H4K5/8/20-triacetyl peptides did result in increased NanoBRET signal with EC_50_ values of 7.5 ± 3.5 *µ*M and 0.98 ± 0.23 *µ*M, respectively (Figure 4A). The NanoBRET signal increased in proportion to the distance between H4K5/8-diacetyl and the third acetyl-lysine residue, likely because greater spacing allows for higher bivalent occupancy (Figure 4A). It is also possible that the increased NanoBRET signal arises from simultaneous binding of one Brd4 protein to H4K4/8-diacetyl and another Brd4 protein to either H4K16ac or H4K20ac on the same peptide. In either case, the ability of either one or two Brd4 proteins to simultaneously bind multiple acetyl-lysine residues on the same peptide does not translate into a biologically relevant increase in peptide binding affinity as all three triacetylated peptides bound to the Brd4 tandem bromodomains with approximately the same affinity (*K*_d_ = 17-24 *µ*M) as determined by ITC (Figure 4C-E). Instead, we hypothesize the 3.2-to 4.5-fold tighter Brd4 tandem bromodomain affinity toward triacetylated peptides relative to the diacetylated peptide (*K*_d_ = 76 *µ*M) is due to the increased avidity of multiple acetyl-lysine binding sites available to individual Brd4 bromodomains on a histone H4 tail (Figure 4B-E, Table S2). As a result, we hypothesize bivalent engagement of multiply acetylated histone H4 tails is not critical for Brd4 regulation of transcription.

**Fig. 4.**
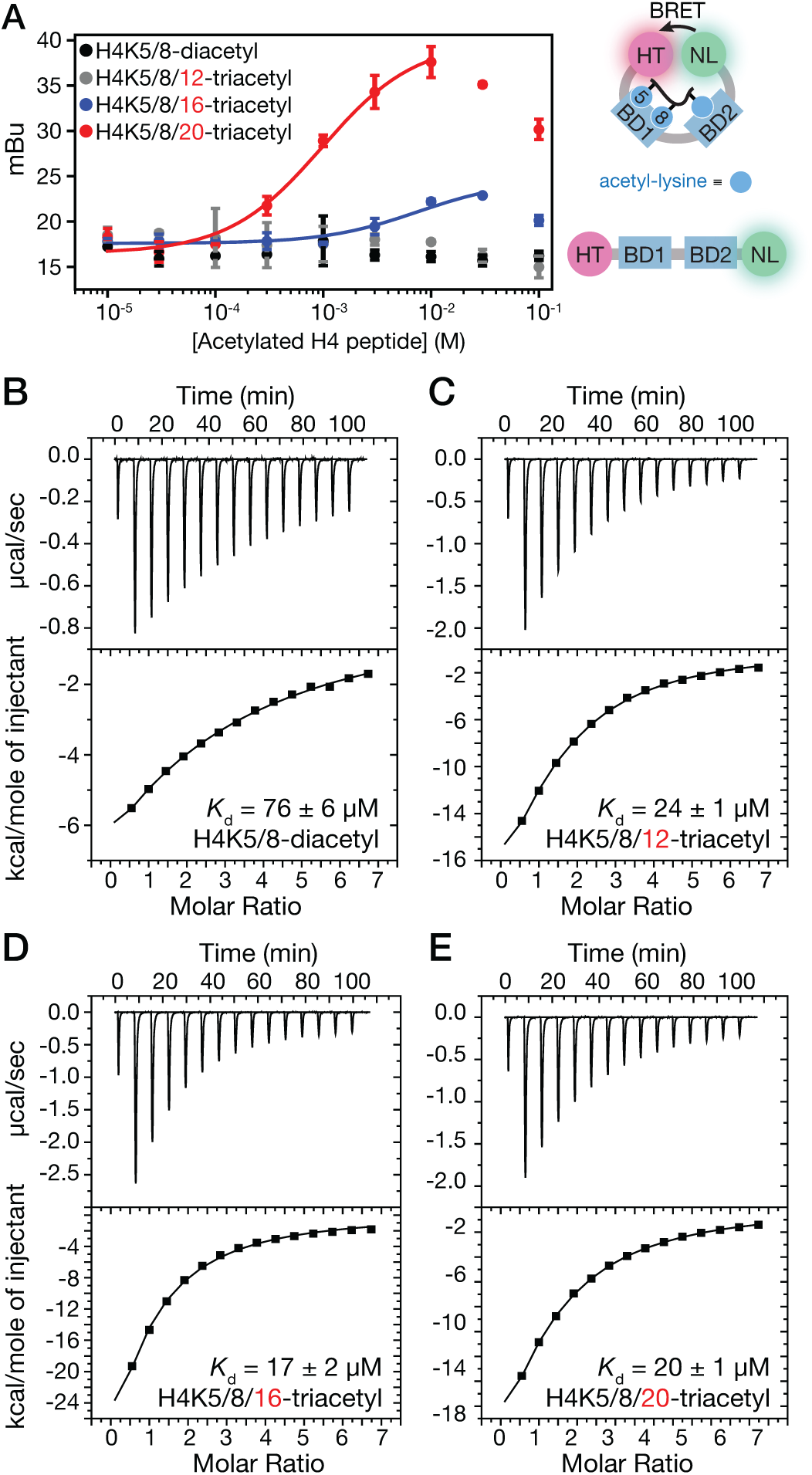
Bivalent tandem bromodomain engagement of acetyl-lysine residues is not responsible for tighter Brd4 binding affinity toward triversus di-acetylated histone H4 tails. (A) NanoBRET signal arising from HaloTag-Brd4-BD1_BD2-NanoLuc titration with histone H4K5/8-diacetyl (black), H4K5/8/12-triacetyl (grey), H4K5/8/16-triacetyl (blue) and H4K5/8/20-triacetyl peptides (red). Representative ITC traces of Brd4 tandem bromodomain (aa 36-460) binding to histone (B) H4K5/8-diacetyl (C) H4K5/8/12-triacetyl (D) H4K5/8/16-triacetyl and (E) H4K5/8/20-acetyl peptides.

### Implications of Brd4-mediated scaffolding of acetylated nucleosomes on 3D chromatin organization

Previous high-resolution Hi-C chromatin conformation capture experiments revealed chromatin 3D structure is partitioned into compartmental domains ranging from 40 kb to 3 Mb in size that correlate with the chromatin transcriptional state (61, 62). Since tandem Brd4 bromodomains scaffold acetylated nucleosomes *in vitro* and interaction of the Brd4 CTD with the RNAP II CTD is crucial for initiation of transcription elongation (63), we hypothesize that the tandem *N*-terminal bromodomains scaffold acetylated nucleosomes to help assemble and maintain 3D chromatin architecture within transcriptionally active compartmental domains. To test this hypothesis, compartmental domains annotated by Rao *et al.* (61) (GSE63525) were separated into transcriptionally active and inactive groups based on ChIP-seq signal for H3K27ac (active) and H3K4me1 (inactive) (61) using k-means clustering (Figure S6). Consistent with our hypothesis that Brd4 cooperates with RNAP II within compartmental domains, Brd4 ChIP-seq signal was preferentially enriched throughout transcriptionally active relative to inactive compartmental domains across multiple cell-lines for which publicly available data sets exist, including IMR90 (*p* < 10^-165^), HUVEC (*p* < 10^-154^), and K562 (*p* < 10^-121^) cells (Figure 5A).

**Fig. 5.**
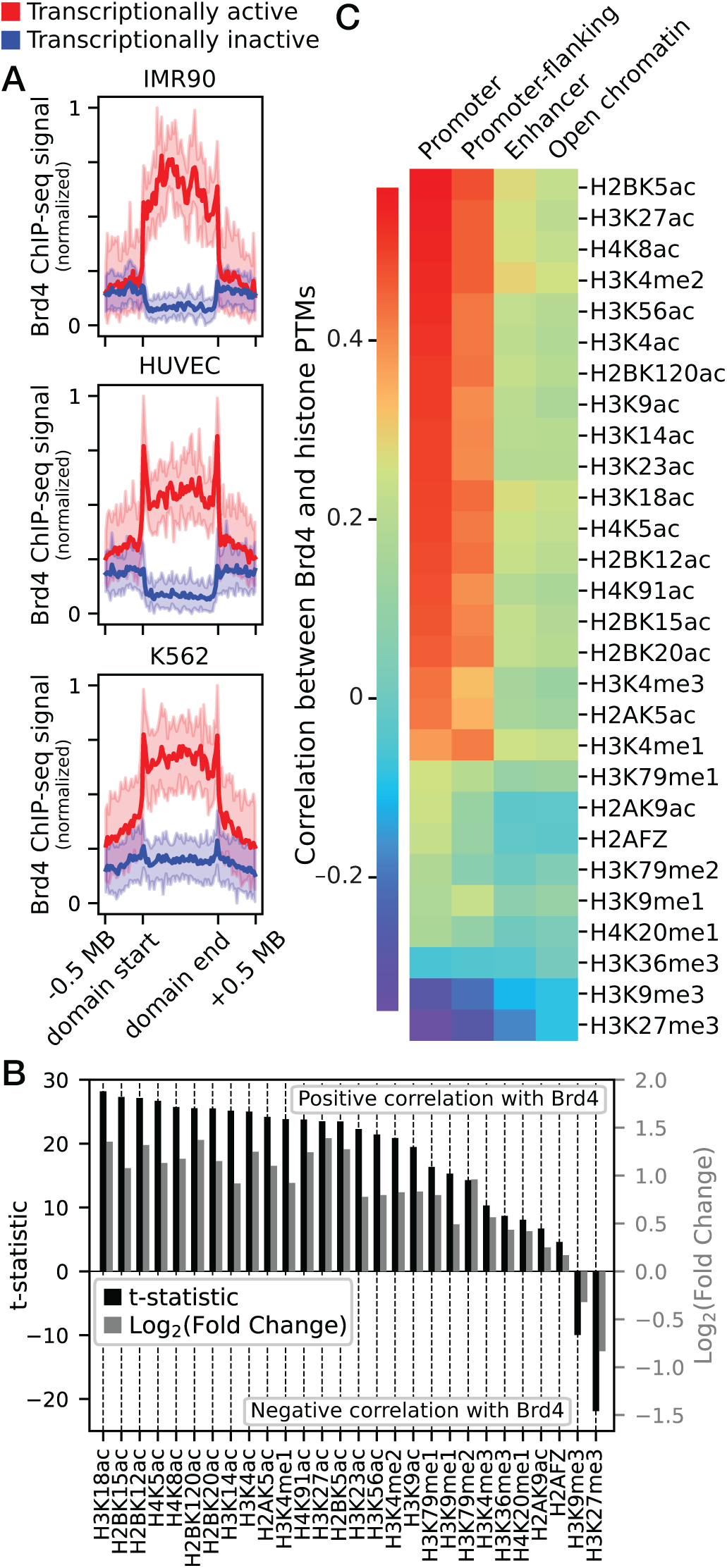
Association between Brd4 and histone acetylation across compartmental domains may contribute to higher-order chromatin structure. (A) Normalized Brd4 ChIP-seq profiles are enriched across transcriptionally active (red) relative to inactive (blue) compartmental domains in IMR90, HUVEC, and K562 cell lines. (B) Bar graph showing significance (two-sided t-statistic, black) and mean fold change (grey) of epigenetic histone PTM ChIP-seq signal over high-versus low-Brd4 occupancy compartmental domains in IMR90 cells. Brd4 demonstrates more significant association with histone lysine acetylation in comparison to non-acetyl histone PTMs. (C) Spearman correlations between ChIP-seq signal for Brd4 and histone PTMs at Ensemble regulatory regions within compartmental domains. Brd4 association with histone lysine acetylation is highest at promoters and promoter-flanking regions.

Histone PTMs have previously been used to identify and classify compartmental domains (62, 64, 65). However, only a subset of known histone modifications were analyzed in these studies including histone H2A family member Z (H2AFZ), H3K27, H3K9 and H4K16 acetylation, as well as H3K27, H3K36, H3K4, H3K79 and H4K20 methylation. Importantly, the Brd4 bromodomains do not bind H3K27, H3K9 or H4K16 acetylation with high affinity *in vitro* (13, 33, 34) suggesting that these acetylation sites do not directly recruit Brd4 to transcriptionally active compartmental domains. To identify histone acetylation sites that correlate with Brd4 occupancy across compartmental domains, we exploited ChIP-seq datasets for 28 distinct histone PTMs (including 17 histone acetylation sites) deposited in the Encyclopedia of DNA Elements (ENCODE) dataset (66). ChIP-seq signals for each histone modification were first binned across IMR90 transcriptionally active and inactive compartmental domains and calculated mean signal per base-pair over each domain. After separating IMR90 compartmental domains based on high or low Brd4 ChIP-seq signal using k-means clustering (Figure S7), high Brd4 occupancy compartmental domains were found to be selectively enriched for histone acetylation over histone methylation and H2AFZ relative to low Brd4 occupancy compartmental domains (Figure 5B). We and others have shown that the tightest affinity histone-Brd4 binding interaction occurs between Brd4-BD1 and histone H4 tails that are diacetylated at lysine residues 5 and 8 (13, 33, 34). Accordingly, H4K5ac and H4K8ac were among the most significantly enriched (*p* < 10^-124^ and *p* < 10^-117^, respectively) histone modifications across high Brd4 occupancy relative to low Brd4 occupancy compartmental domains. Also, H3K18ac and H3K14ac showed the most significant co-enrichment (*p* < 10^-135^ and *p* < 10^-114^, respectively) with Brd4 among histone H3 PTMs. These results are consistent with previous *in vitro* binding assays (13) and cell-based pull-downs (67) that demonstrate that H3K14ac and H3K18ac, in addition to H4K5ac, are among the tightest affinity histone binding targets of Brd4-BD2.

To determine the types of regulatory chromatin regions that may participate in acetylation-dependent scaffolding by the Brd4 tandem bromodomains, compartmental domains were separated into promoter, promoter-flanking, enhancer, and open chromatin regions using Ensembl regulatory element annotations (68). Brd4 correlation with histone acetylation was greatest at promoters and promoter-flanking regions compared to enhancers and open chromatin regions (Figure 5C). In addition, H4K5ac, H5K8ac, H3K14ac and H3K18ac were among the highest Brd4-correlating PTMs at promoters (Spearman correlation ≥ 0.4). These findings suggest a potential role for the Brd4 tandem bromodomains in maintaining promoter-promoter interactions, consistent with facilitating transcriptional co-regulation of interacting gene networks (69).

## Discussion

BET proteins bind to acetylated chromatin via their *N*-terminal bromodomains and recruit components of the transcription machinery via an extra-terminal (ET) protein-protein interaction domain. BrdT and Brd4 also include a conserved *C*-terminal domain (CTD) that interacts with the positive transcription elongation factor b (P-TEFb) and the RNAP II CTD during activation of transcription elongation (70–72). Whereas expression of BrdT is limited to the testes and ovaries among adult tissues, Brd4 is ubiquitously expressed (15) and widely implicated in human disease (16). Despite the biological significance of Brd4-mediated transcription regulation, it is unclear why two bromodomains in tandem are necessary for chromatin binding and full Brd4 activity. Here, Brd4 was used to investigate potential models for multivalent binding of BET tandem bromodomains to acetylated chromatin.

In our SAXS analyses, BrdT, Brd2, Brd3 and Brd4 all demonstrated similar *R*_*g*_ (48.6-55.4 Å) and *D*_*max*_ (181-200 Å) values (Figure 2A-B and S2, Table S1) consistent with previous SAXS measurements (73). Consequently, the high degree of inter-bromodomain flexibility demonstrated by our SAXS-guided Rosetta *ab initio* modeling of the Brd4 linker sequence is likely conserved across all four BET proteins. In contrast, a crystal structure (PDB ID: 3UV5) (13) of the Taf1 tandem bromodomains (*R*_*g*_ = 30.5 ± 0.5 Å and *D*_*max*_ < 103 ± 5 Å) displays an inter-bromodomain acetyl-lysine binding site distance of 60.1 Å. Furthermore, the majority of the remaining human tandem bromodomain-containing proteins contain short inter-bromodomain linkers relative to BET tandem bromodomains, including Brwd1 (81 aa, UniProt ID: Q9NSI6), Brwd3 (87 aa, Q6RI45), Pbrm1 (64, 128, 66, 66 and 44 aa for BD1-BD2, BD2-BD3, BD3-BD4, BD4-BD5 and BD5-BD6, respectively, Q86U86), PHIP (85 aa, Q8WWQ0) and Taf1L (51 aa, Q8IZX4). The only exception is Brd8 (Q9H0E9) which contains a 324 aa inter-bromodomain linker. As a result, we hypothesize BET tandem bromodomains have greater conformational flexibility compared to other human multiple bromodomain-containing proteins (with the possible exception of Brd8), perhaps allowing for unique and diverse acetyl-lysine-dependent epigenetic functions to be carried out by BET proteins.

Our SAXS-guided Rosetta modeling indicated that the Brd4 tandem bromodomains may access conformations in which the two acetyl-lysine binding sites reside as little as 15 Å apart, perhaps allowing for bivalent binding toward adjacent acetylation sites on individual histone tails. These results are consistent with the known ability of bivalent BET bromodomain inhibitors to simultaneously engage both acetyl-lysine binding sites (59, 74, 75). Previous studies of the Taf1 tandem bromodomains demonstrated 8-to 29-fold tighter binding affinity toward diacetylated (*K*_d_ = 1.4 to 5.3 *µ*M) relative to monoacetylated histone H4 tail mimic peptides (*K*_d_ ∼40 *µ*M) and proposed a model in which the two Taf1 bromodomains bivalently engage separate acetyl-lysine residues on a single peptide. In our studies, Brd4 binding was also enhanced in ITC affinity measurements (3.2 to 4.5-fold, Figure 4B-E) by increasing the number of acetyl-lysine residues on histone H4 tail mimic peptides from H4K5/8-diacetyl to H4K5/8/12-triacetyl, H4K5/8/16-triacetyl or H4K5/8/20-triacetyl. However, only histone H4K5/8/16-triacetyl and H4K5/8/20-triacetyl peptides and not the H4K5/8/12-triacetyl peptide showed evidence of simultaneous binding to two separate bromodomains in our NanoBRET assay (Figure 4A). Therefore, the correlation between occupancy of acetylated histone H4 tail residues and Brd4 binding affinity likely results from the increased number of acetyl-lysine binding sites available for individual bromodomains, not bivalent engagement of a single histone tail peptide by a single tandem bromodomain construct (Figure 1B).

The significance of triacetylated histone H4 tails in improving Brd4 tandem bromodomain affinity may be further limited in the context of a nucleosome. For instance, previous studies by Miller *et al.* showed that BrdT-BD2 binds H3K18/23-diacetyl histone tail peptides but not nucleosomes harboring these PTMs (76). Therefore, while Brd4-BD1 binds to H4K5/8-diacetyl peptides and nucleosomes with equal affinity (76) and our NanoBRET assays indicate the Brd4 tandem bromodomains can also simultaneously bind H4K16ac and H4K20ac on histone tail peptides, it is questionable whether these sites are recognized by Brd4 in the context of nucleosomes. As a result, while Brd4 tandem bromodomains are capable of simultaneously binding at least two distinct acetylated regions on single H4 tail peptides (Figure 1B), this activity is likely not central to their biological function.

The relatively long and flexible BET inter-bromodomain linkers compared to non-BET tandem bromdomains suggests that BET tandem bromodomains have evolved to simultaneously engage two separate acetylated proteins. The discovery of acetylated non-histone BET bromodomain binding proteins has led others to propose that the BET tandem bromodomains recruit transcription factors to chromatin regions via bivalent acetyl-lysine recognition (Figure 1A). For instance, Brd4 binds acetylated forms of cyclin T1 (55, 72), the RelA subunit of NF-*κ*B (77, 78), and the transcription factor Twist (79). In addition, a recent study by Lambert *et al.* (73) suggests BET protein interactions are more widespread than previously appreciated and demonstrated structural evidence of Brd4 bromodomain binding to numerous acetylated nuclear proteins. However, the ability of tandem bromodomains to simultaneously engage separate acetylated proteins had not been tested directly. Using sucrose gradient binding assays (Figure 3), we demonstrated for the first time that the Brd4 tandem bromodomains can scaffold separate acetylated nucleosomes, bringing them into close proximity (Figure 1A).

Previous studies indicate bromodomain-nucleosome association may in some cases be driven by interactions other than acetyl-lysine recognition. The yeast BET protein homologs, Bromodomain factor 1 and 2 (Bdf1 and Bdf2) demonstrate weak affinity for unacetylated histone H4 tails, likely due to favorable electrostatic contacts (36). A subset of human bromodomains also bind nonspecifically to DNA to enhance (76) or even drive (80) association with acetylated nucleosomes. For instance, previous *in vitro* studies indicated that the DNA-binding activity of BrdT-BD1 is not sufficient for interaction with nucleosomes and only serves to strengthen the affinity of acetylation-dependent nucleosome interactions (76), whereas binding of the Brg1 bromodomain to nucleosomes is primarily driven by DNA interactions (80). Brd4-BD2 was also previously shown to bind DNA in electrophoretic mobility shift assays (76). In our studies, while interactions with nucleosomal DNA and residues adjacent to acetyl-lysine on histone tails may contribute to Brd4 association, only bromodomain acetyl-lysine binding was necessary for nucleosome scaffolding activity.

Bromodomain-mediated nucleosome scaffolding presents a potential mechanism for dynamic control of chromatin 3D structure via reversible epigenetic lysine acetylation. Indeed, differences in higher-order structural involvement across chromatin regions are associated with distinct histone PTM patterns (61). In general, 3D chromatin organization is defined by combinations of cohesin-mediated CCCTC-binding factor (CTCF) loops and compartmental domains that are not associated with CTCF peaks (61). While CTCF loops are formed through CTCF- and cohesion-mediated loop extrusion (81–85), formation and maintenance of compartmental domains is associated with chromatin transcriptional state (62, 64, 65, 86). Recent studies proposed a ‘transcription factory’ hypothesis in which cooperative interactions between multivalent transcription factors bound to RNAP II drive transcriptionally-active compartmental-domain formation (87, 88). However, the multivalent chromatin-binding proteins involved in nucleosome scaffolding within compartmental domains are poorly defined. Previous studies showed that loss of Brd4 results in global chromatin decondensation in human cell lines and that the Brd4 *C*-terminal domain (CTD) in addition to the tandem bromodomains are necessary to maintain normal chromatin structure (44). As a result, Brd4 represents an intriguing candidate for multivalently scaffolding acetylated chromatin in the context of transcription factories.

Through bioinformatic analyses of publicly available ChIP-seq datasets deposited in the GEO and ENCODE databases, we found that Brd4 is enriched throughout transcriptionally active and not transcriptionally inactive compartmental domains in diverse cell types (Figure 5A). Brd4 occupancy across compartmental domains correlated strongly with histone acetylation sites that are known peptide binding targets of the Brd4 bromodomains, including H3K14ac, H3K18ac, H4K5ac, and H4K8ac (Figure 5B) (13, 33, 34, 67). In addition to serving as a potential histone tail binding site for Brd4-BD2, Brd4 chromatin binding is proposed to drive acetylation of H3K18 (89, 90), perhaps explaining why this modification is the most significantly enriched PTM at high Brd4 occupancy compartmental domains in our dataset. On the other hand, we found 7 additional histone acetyl-lysine residues (H2AK5ac, H2BK12ac, H2BK15ac, H2BK20ac, H2BK120ac, H3K4ac and H4K91ac) that are not known to interact with Brd4 bromodomains are more significantly (*p* < 10^-102^) co-enriched with Brd4 across compartmental domains than the prototypical transcriptionally activating H3K4me1 and H3K27ac (*p* < 10^-103^ and *p* < 10^-100^ respectively) histone PTMs. Since levels of each of these histone acetyl-lysines correlate strongly (Spearman correlation coefficient ≥ 0.90) with H4K5ac, H4K8ac, H3K14ac and H3K18ac across compartmental domains (Figure S8), these PTMs may be upregulated by epigenetic writer and eraser enzymes present in transcription factor complexes that involve Brd4 rather than directly recruiting Brd4 to chromatin.

We also found that Brd4 correlates most strongly with histone acetyl-lysine binding targets at promoters relative to other *cis*-regulatory regions (Figure 5C). Previous studies by Li et al. identified ∼20,000 RNAP II-associated intrachromosomal interactions that are conserved across human cell-types and showed that they most commonly involved promoter-promoter (44%) contacts relative to promoter-gene internal (5%), promoter-enhancer (33%), and enhancer-enhancer (20%) contacts (69). Since Brd4 interacts *C*-terminally with the RNAP II CTD, we hypothesize that the tandem bromodomains may function within transcription factories to scaffold acetylated promoters, facilitating co-regulation of transcriptionally related genes. However, this activity would be limited to Brd4 and BrdT, as Brd2 and Brd3 lack the conserved CTD through which Brd4 and BrdT interact with RNAP II (91).

In summary, these analyses suggest a potential role for Brd4 scaffolding acetylated nucleosomes at promoters throughout transcriptionally active chromatin compartments, potentially bringing together transcriptionally co-regulated genes that are distant in primary DNA sequence. Overall, our studies provide direct evidence supporting (internucleosomal) and against (intranucleosomal) different modes of multivalent chromatin engagement by tandem Brd4 bromodomains. Our results provide a mechanistic rationale for investigating the role of Brd4 and other proteins containing linked histone-binding domains in regulating chromatin 3D organization in the cellular context. In addition, there are many proteins encoding linked histone-binding modules beyond the tandem bromodomain-containing proteins (30) that may also be involved in the assembly of higher-order chromatin through histone PTM patterns, which is an interesting avenue for further study.

## Materials and Methods

### SAXS analysis of tandem bromodomain constructs

1, 2, 5 and 10 mg/mL samples of each protein were shipped to SIBYLS (92) overnight. SAXS data were collected via the mail-in program (92, 93) using the SIBYLS beamline 12.3.1 (92) at the Advanced Light Source in Lawrence Berkeley National Laboratory. The 1-dimensional buffer-subtracted SAXS profile at each protein concentration was calculated from the average of 32 measurements using the SIBLYS application, FrameSlice. For each protein sample, the SAXS profiles at different protein concentrations were inspected to exclude contribution from protein aggregation (caused by cumulative radiation damage) before merging to one composite SAXS profile using the SCÅTTER program. Using the SCÅTTER program, Kratky plots, *R*_*g*_ values, maximum dimensions (*D*_max_), and *P*(*r*) functions were calculated.

### Rosetta modeling of the Brd4 inter-bromodomain linker

The input Brd4 tandem bromodomains structure for *ab initio* calculations was prepared by connecting crystal structures of Brd4-BD1 (4KV1, chain A; aa 44-168) and Brd4-BD2 (4KV4, chain A; aa 348-459) with residues 169-348 built in an extended conformation. The side chains of the starting model were prepacked using the Rosetta-fixed backbone design/packing application (using the parameters -ex1, -ex2aro, use_input_sc). The FloppyTail protocol was then used to generate 5000 models of the bromodomain linker in which the complete structure passed *R*_*g*_ and *D*_*max*_ (201 Å) filters determined from the SAXS analyses. The inter-bromodomain distances were then determined by measuring the distance between the conserved Asn residue sidechain amide nitrogen atoms in Brd4-BD1 and BD2 (N140 and N433, respectively) for each model.

### NanoBRET peptide binding assays

Recombinant HaloTag-Brd4-BD1_BD2-NanoLuc (100 nM; expression and purification described in Supporting Information) with (experimental) and without (control) 100 nM HaloTag NanoBRET 618 fluorescent ligand (Promega) was combined with diacetylated or triacetylated histone peptide concentrations ranging from 10 nM to 100 *µ*M in white, flat-bottomed 96-well plates (Corning). Plates were incubated for 30 min at 25 °C before NanoBRET Nano-Glo Substrate (Promega) was added to both control and experimental samples at a final concentration of 10 *µ*M. Plates were read within 10 min using a Tecan Spark plate reader. A corrected BRET ratio was calculated, defined as the ratio of the emission at 610 nm/450 nm for experimental samples (*i.e.*, those treated with HaloTag NanoBRET 618 fluorescent ligand) minus the emission at 610 nm/450 nm for control samples (*i.e.*, those not treated with HaloTag NanoBRET 618 fluorescent ligand). BRET ratios were expressed as milliBRET units (mBU), where 1 mBU corresponds to the corrected BRET ratio multiplied by 1,000.

### Isothermal titration calorimetry

Binding affinities of H4(1-11)K5/8-diacetyl and H4(1-15)K5/8/12-triacetyl, H4(1-19)K5/8/16-triacetyl and H4(1-23)K5/8/20-triacetyl histone peptides toward the Brd4 tandem bromodomains (aa 36-460) were determined using a VP-ITC instrument (MicroCal). Briefly, 0.4 mM H4K5/8diacetyl, H4K5/8/12-triacetyl, H4K5/8/16-triacetyl or H4K5/8/20-triacetyl peptides were injected (1 × 4 *µ*L injection followed by 14 × 16 *µ*L injections) into the cell containing 10 *µ*M Brd4 (aa 36-460), and heats of binding were measured. The buffer used for ITC analysis consisted of 25 mM HEPES (pH 7.5 at 20 °C), 150 mM NaCl and 2% v/v glycerol. Protein concentrations were determined using the method of Bradford (94). The least-squares fits to the binding parameters Δ*H* °, *K*_d_, and *N* were determined from the raw data using Origin (OriginLab).

### Nucleosome purification

Mono-nucleosomes were purified as previously described (34). Briefly, 100 *µ*L of chick erythrocyte or calf thymus nuclei (10 mg/mL) in 0.25 M sucrose, 10 mM MgCl_2_ and 10 mM Tris (pH 8.0) were added to 200 *µ*L of 100 mM NaCl, 1 mM CaCl_2_ and 40 mM Tris (pH 8.0). After 3 min equilibration at 35 °C, 2 *µ*L of micrococcal nuclease (5 U/mL) was added and the solution was incubated at 35 °C for 12.5 min. The nuclease reaction was then quenched with 6 *µ*L of 250 mM EDTA and the mixture was pelleted for 4 min at 13,200 rpm. The pellet was resuspended in 300 *µ*L of 1 mM EDTA and pelleted for 4 min at 13,200 rpm and 200 *µ*L of the resulting supernatant was applied to a 4 mL sucrose gradient (5–20% w/v sucrose with 1 mM EDTA, pH 8.0) and centrifuged at 55,000 × g for 3.5 h. The sucrose gradient was then collected in fractions, nucleoproteins were digested with Proteinase K and the DNA analyzed by agarose (1.5% w/v) gel electrophoresis. Concentrations of purified nucleosomes were determined according to DNA absorbance at 260 nm using an extinction coefficient of 6,600 M^-1^cm^-1^.

### Sucrose gradient binding assay

500 nM mono-nucleosomes were combined with 10 *µ*M recombinantly purified His_6_-tagged Brd4 (aa 36-460) WT, N140F or N433F in 25 mM HEPES (pH 7.5 at 20 °C) and 150 mM NaCl. Each sample was incubated for 30 min at 25 °C and applied to a 4 mL sucrose gradient (5–20% w/v sucrose with 1 mM EDTA, pH 8.0) and centrifuged at 55,000 × g for 3.5 h. The sucrose gradient was then collected in fractions. Nucleosome-containing fractions were identified by agarose gel electrophoresis combined with ethidium bromide staining as described above and Brd4-containing fractions were identified by anti-His_6_ tag immunoblotting. Nitrocellulose membranes were first blocked using PBST with 3% w/v BSA. The blocked membranes were incubated with an anti-His_6_ tag primary antibody (Abgent, AM1010A) at a dilution of 1:1000 in PBST with 1.5% w/v BSA overnight at 4 °C. Membranes were then washed 3×5 min with PBST and incubated with a 1:10,000 dilution of Anti-mouse IgG secondary antibody HRP (GeneTex, GTX213111-01) in PBST with 1.5% w/v BSA for 1 h at room temperature. Finally, membranes were washed 3×5 min with PBST and chemiluminescence was detected.

### ChIP-Seq data analysis

BED files containing compartmental domain annotations in HUVEC, K562 and IMR90 cells were obtained as text files from GSE63525 (61). Brd4 ChIP-Seq datasets from HUVEC (GSM1305201) (49), K562 (ENCFF260JHC) and IMR90 (GSM1915116) (95) cells were converted from Bedgraph and Wig to BigWig format when necessary using the UCSC Genome Browser applications bedGraphToBigWig and wigToBigWig, respectively. HUVEC ChIP-Seq data was converted from the hg18 to hg19 genome assembly using the CrossMap Python package. Histone ChIP-Seq data was obtained from ENCODE reference epigenome datasets for HUVEC (ENCSR194DQD), K562 (ENCSR612NLL) and IMR90 (ENCSR596VTT) cells as BigWig files consisting fold-change signal over control from two merged replicates. Bigwig files were binned over compartmental domains using the DeepTools computeMatrix function in scale-regions mode with region body length of 1 Mbp and upstream and downstream distances of 0.5 Mbp. Compartmental domains were clustered based on active enhancer modification (H3K27ac and H3K4me1) or Brd4 ChIP-Seq signal by passing the kmeans flag to the DeepTools plotProfile function. IMR90 ChIP-Seq signal across high and low Brd4 occupancy compartmental domain clusters were calculated from the mean signal per base-pair over the entire range of each domain and significance was calculated from pooled mean signals using the ttest_ind function from the Scipy Python package assuming unequal sample variance. Spearman correlation coefficients between ChIP-seq signal for Brd4 and IMR90 histone marks over Ensembl regulatory features and between mean IMR90 histone mark signal per base pair over compartmental domains were calculated using the Scipy spearmanr function.

## ACKNOWLEDGMENTS

We thank R.B. Hill for his detailed comments. This work was supported in part by NIH grant R35 GM128840 (to B.C.S.), American Heart Association grant 15SDG25830057 (to B.C.S.), Institutional Research Grants 14-247-29-IRG and 86-004-26-IRG from the American Cancer Society to B. C. S.), American Diabetes Association grant 1-18-IBS-068 (to B.C.S.), the Michael Keelan, Jr., MD, Research Foundation (to B.C.S.), and the Advancing a Healthier Wisconsin endowment (to B.C.S.). SAXS data collection was conducted at the Advanced Light Source (ALS), a national user facility operated by Lawrence Berkeley National Laboratory on behalf of the Department of Energy, Office of Basic Energy Sciences, through the Integrated Diffraction Analysis Technologies (IDAT) program, supported by DOE Office of Biological and Environmental Research. Additional support comes from the National Institute of Health project ALS-ENABLE (P30 GM124169) and a High-End Instrumentation Grant S10ODO18413. Computational resources were provided by the Research and Computing Center of the Medical College of Wisconsin. M.D.O. is a member of the Medical Scientist Training Program at Medical College of Wisconsin, which is supported in part by National Institutes of Health Training Grant T32-GM080202 from NIGMS.

