## Supporting Information for "Nucleosome scaffolding by Brd4 tandem bromodomains in acetylation-dependent chromatin compartmentalization"

**This PDF file includes:**

Supplementary text
Figs. S1 to S8
Tables S1 to S2

### Supporting Information Text

#### Methods.

**Protein expression and purification.** The tandem bromodomains of human Brd4 (aa 36-460), Brd3 (aa 25-416) and Brd2 (aa 71-455) and BrdT (aa 18-383) were subcloned (Genscript) between the NdeI and XhoI restriction sites of a pET28b vector modified to replace the thrombin cleavage site with a tobacco etch virus (TEV) protease cleavage site between the *N*-terminal $His_6$ tag and the BET bromodomain protein coding region. Brd4 Asn mutants (N140F and N433F) were generated using site directed mutagenesis (Genscript). The $His_6$-tagged TAF1 tandem bromodomain construct (aa 1373-1635) in the pNIC28-Bsa4 vector was a gift from Nicola Burgess-Brown (Addgene plasmid #39118; http://n2t.net/addgene:39118 ; RRID:Addgene_39118). HaloTag-BRD4-BD1__BD2-NanoLuc (aa 44-460) was generated (Genscript) by subcloning NanoLuc-Brd4 (aa 44-460) from NL-Brd4 pFC27K vector (Promega) into the pH6HTC $His_6$HaloTag T7 vector (Promega). The recombinant $His_6$-tagged tandem bromodomain constructs were purified from BL21(DE3) *E. coli* using nickel affinity chromatography. Cells were transformed and grown in 2–4 L of LB in the presence of 50 $\mu$g/mL kanamycin or 100 $\mu$g/mL ampicillin to an optical density of 0.6–0.8 at 600 nm. Protein expression was induced with 0.1 mM IPTG and the cells were incubated overnight at 18 °C. Cells from each 1 L culture were harvested by centrifugation at 5,000 × *g* and re-suspended in 30 mL of lysis buffer (50 mM HEPES pH 7.5 at 20 °C, 500 mM NaCl, 5% v/v glycerol and 2.5 mM imidazole) supplemented with protease inhibitors (0.3 $\mu$M aprotinin, 1 $\mu$M E-64, 1 $\mu$M leupeptin, 1 $\mu$M bestatin, 1 $\mu$M pepstatin and 100 $\mu$M PMSF). Cells were lysed by sonication and lysates were cleared by centrifugation for 30 min at 30,000 × *g*. The lysate supernatents were then applied to Ni-NTA resin (0.75 mL resin/L of bacterial culture) and rocked for 1 h at 4 °C. The supernatant was discarded and the Ni-NTA resin was applied to a column and washed twice with 25 mL of lysis buffer. The protein was eluted using a step gradient of increasing concentrations of imidazole in lysis buffer (5 mL of 50, 100, 150, 200 and 250 mM imidazole). Fractions were monitored by SDS-PAGE and those containing recombinant protein were concentrated to a volume of 1 mL and applied to a HiLoad 16/600 Superdex 75 pg column (Bio-Rad) and eluted with 25 mM HEPES (pH 7.5 at 20 °C), 150 mM NaCl, and 2% v/v glycerol. For EPR and SAXS studies, $His_6$ Ni-NTA affinity tags were removed by Tobacco Etch Virus (TEV) protease cleavage prior to gel filtration. Samples containing recombinant protein were identified by SDS-PAGE and concentrated to 5–10 mg/mL, flash frozen in liquid nitrogen and stored at −80 °C until used.

**Synthesis of JQ1-TEMPO EPR probe.** JQ1 (5.0 mg, 12.7 $\mu$mol) was dissolved in 1 mL of DCM, 0.25 mL of TFA was added and the reaction was stirred overnight at 25 °C to cleave the tBu ester to the free carboxylic acid. The reaction was dried under reduced pressure then redissolved and dried three times with $\tilde{1}$ mL chloroform to remove residual formic acid. The resulting solid was dissolved in 156.7 $\mu$L anhydrous DMF and to the solution was added HBTU (6.22 mg, 16.4 $\mu$mol), DIEA (7.1 $\mu$L, 54.7 $\mu$mol) and 4-Amino-TEMPO (2.8 mg, 16.4 $\mu$mol). The reaction was stirred at 25 °C for 30 h and the resulting JQ1-TEMPO was purified by semipreparative HPLC on a 5 $\mu$m particle size Hypersil GOLD C18 column (ThermoFisher, 4.6 × 250 mm) using an Agilent 1100 series HPLC eluting with a gradient of 5–95% v/v acetonitrile in water with 0.1% v/v TFA. The mass of JQ1-TEMPO was confirmed by direct injection ESI mass spectrometry (QExactive, Thermo Scientific). HRMS (ESI): Exact mass calculated for $C_{28}H_{35}ClN_6O_2S$ $[M+H]^+$ 554.2231, found 554.2242.

**Electron paramagnetic resonance.** 75 $\mu$M Brd4 (aa 36-460) in 25 mM HEPES, 150 mM NaCl, 25% v/v glycerol was titrated with JQ1-TEMPO to 350 $\mu$M where saturation was reached in the continuous wave (X-band) spectrum and double electron-electron resonance measurements were subsequently recorded.

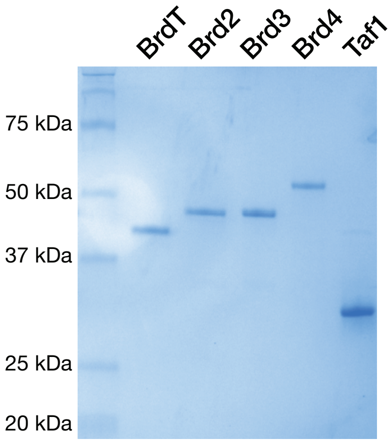


**Fig. S1.** Identity and purity of recombinant tandem bromodomain proteins. SDS-PAGE followed by Coomassie staining of 1 µg each of $His_6$-tagged BrdT (44.6 kDa), Brd2 (45.9 kDa), Brd3 (46.2 kDa), Brd4 (50.5 kDa) and Taf1 (34.6 kDa).

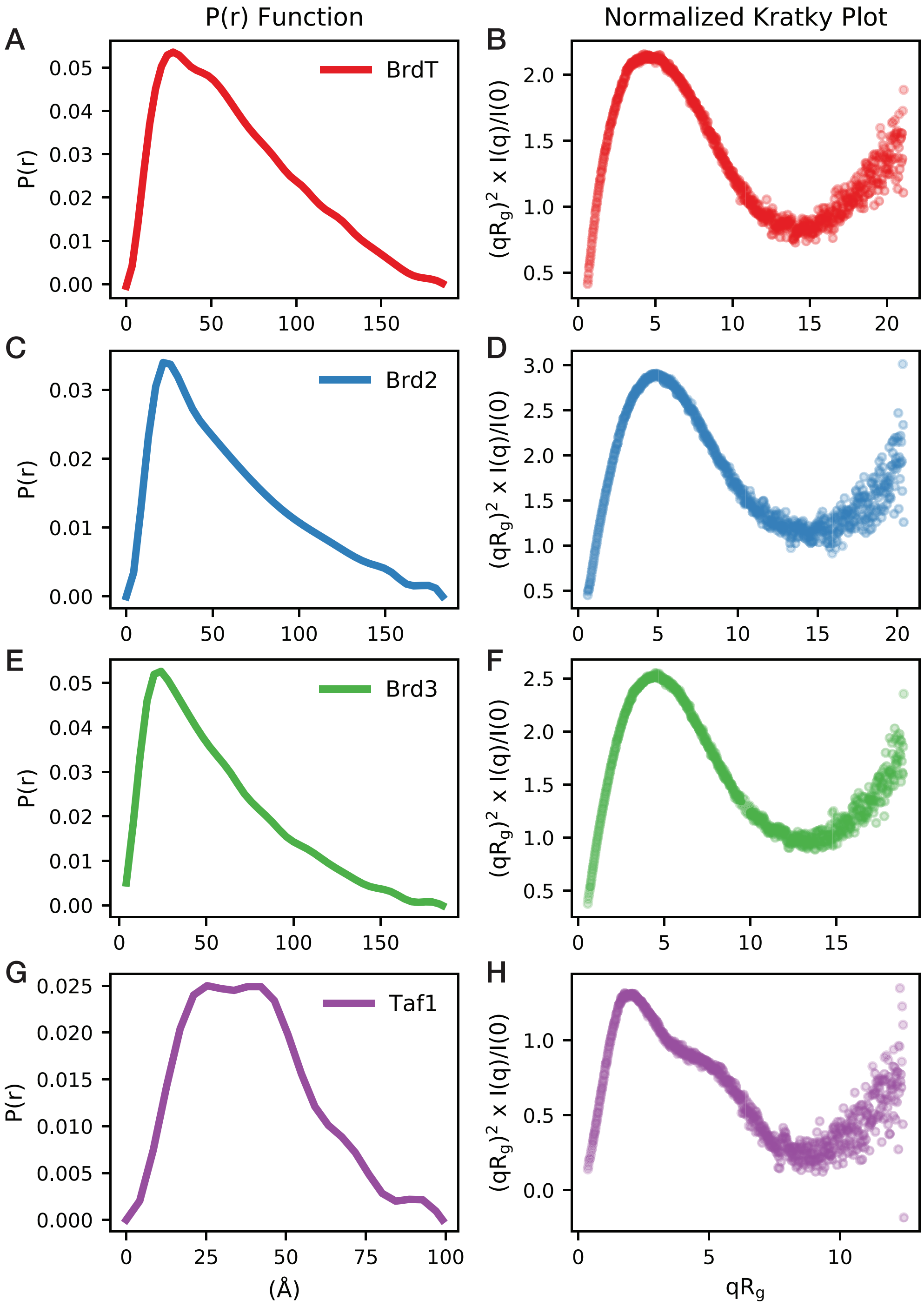


**Fig. S2.** SAXS analysis of tandem bromodomains other than the Brd4 bromodomains. *P(r)* distributions and $R_g$-normalized Kratky representations for (A-B) BrdT, (C-D) Brd2, (E-F) Brd3 and (G-H) Taf1. SAXS analysis reveals relatively long and cylindrical shapes for BET tandem bromodomains compared to the relatively globular Taf1 tandem bromodomains.

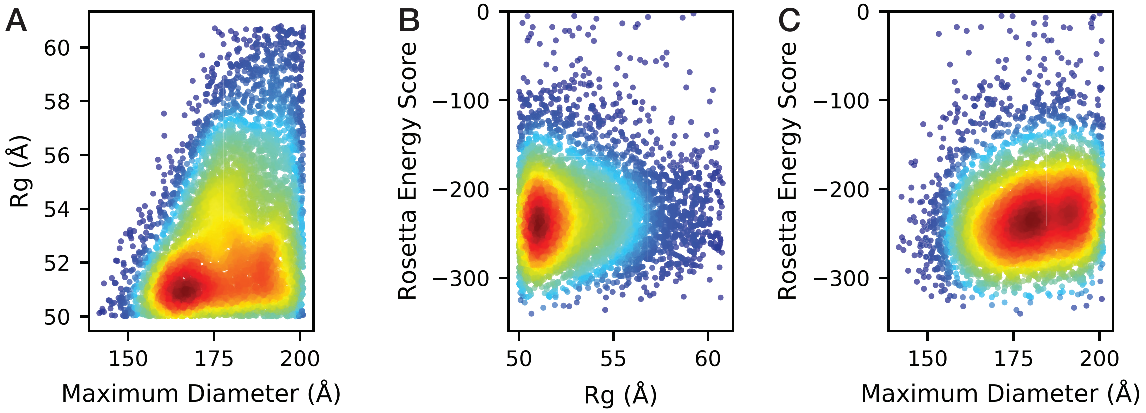


**Fig. S3.** Filtering of Rosetta Brd4 tandem bromodomain models by SAXS constraints. (A) Rosetta models were constrained to $D_{max} \leq 201$ Å and $49.9 \leq R_g \leq 59.9$. Rosetta energy score convergence was not observed across allowed (B) $R_g$ or (C) $D_{max}$ ranges.

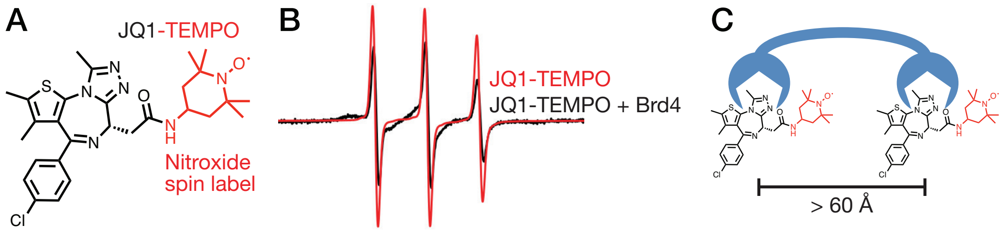


**Fig. S4.** EPR-based strategy for Brd4 tandem bromodomain inter-acetyl-lysine binding site measurements. (A) Structure of JQ1-TEMPO EPR probe. (B) Continuous wave (CW) EPR detection of JQ1-TEMPO binding to the Brd4 tandem bromodomains. (C) No double electron-electron resonance (DEER) signal was observed in the JQ1-TEMPO + Brd4 sample, indicating the two bromodomain acetyl-lysine binding pockets exist predominantly > 60 Å apart in solution.

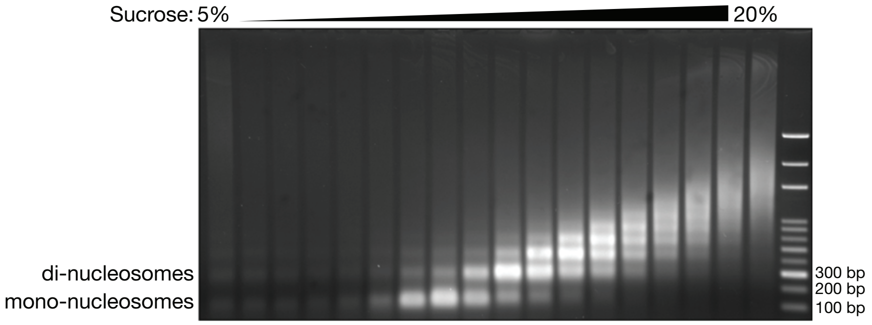


**Fig. S5.** Representative sucrose gradient purification of calf thymus nucleosomes. Fractions containing primarily mono-nucleosomes were combined and used in subsequent Brd4 binding assays.

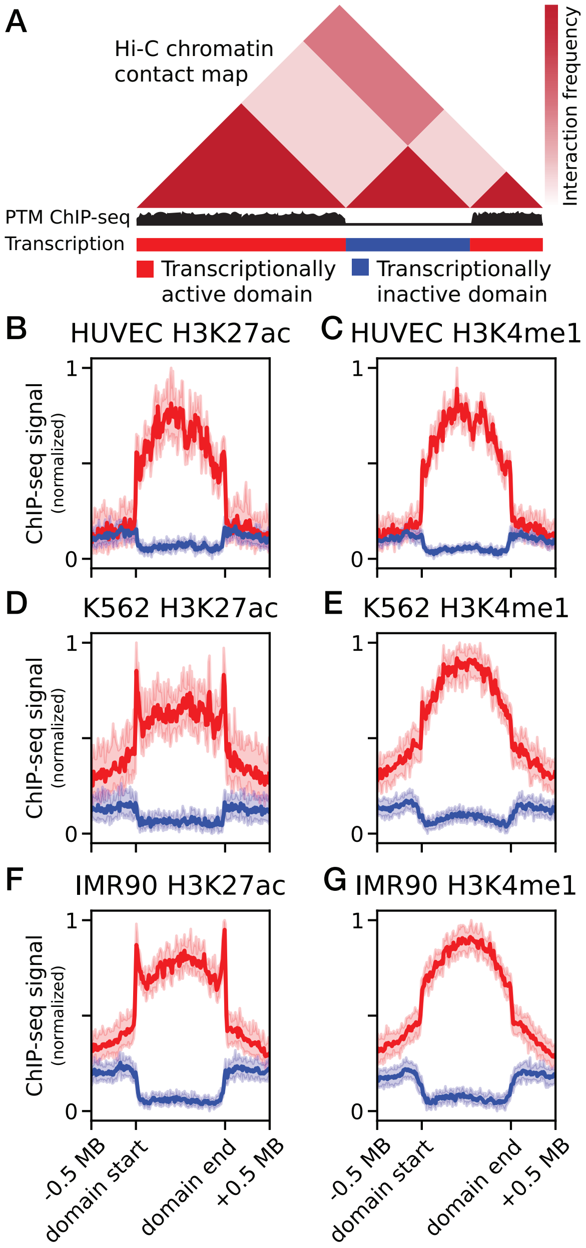


**Fig. S6.** K-means clustering of H3K27ac and H3K4me1 ChIP-seq profiles across annotated compartmental domains (GSE63525). (A) Schematic depicting transcriptionally active and inactive compartmental domain designations based on H3K27ac and H3K4me1 ChIP-seq signal clustering. Average H3K27ac and H3K4me1 ChIP-seq signals are plotted accross compartmental for (B-C) IMR90, (D-E) HUVEC, and (F-G) K562 cell lines. Clusters containing high (red) and low (blue) H3K27ac/H3K4me1 occupancy were used to define transcriptionally active and inactive compartmental domains, respectively.

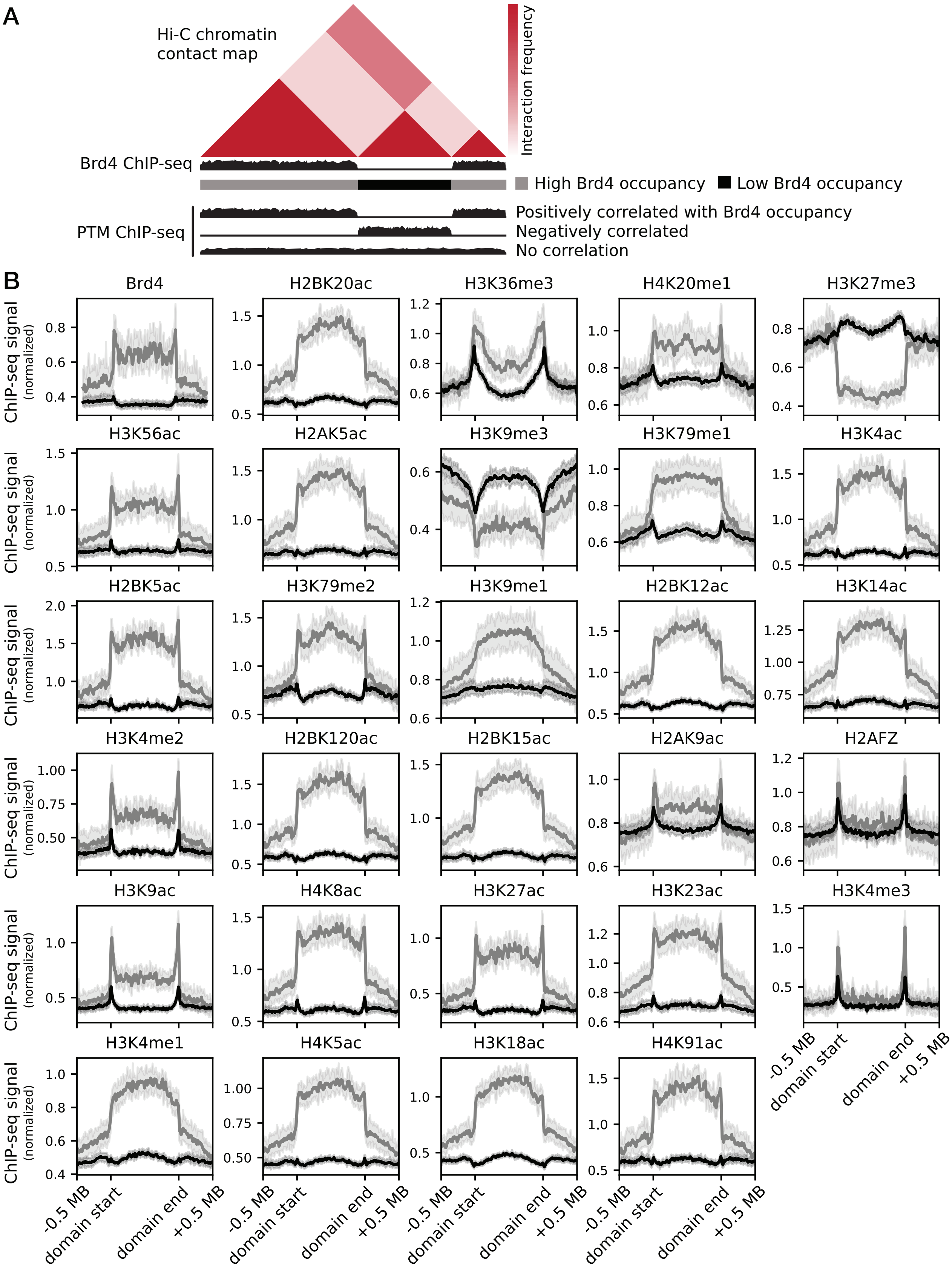


**Fig. S7.** Association between Brd4 ChIP-seq signal and epigenetic histone PTMs across IMR90 compartmental domains. (B) IMR90 compartmental domains were clustered into high (gray) and low (black) Brd4-occupancy (upper left corner) subsets using the K-means algorithm. ChIP-seq profiles for 28 histone post-translational modifications were calculated across compartmental domains in the high (gray) and low (black) Brd4 occupancy subsets to identify significant differences (two-tailed t-test allowing for unequal population variance) and mean fold-changes in histone PTM signals.

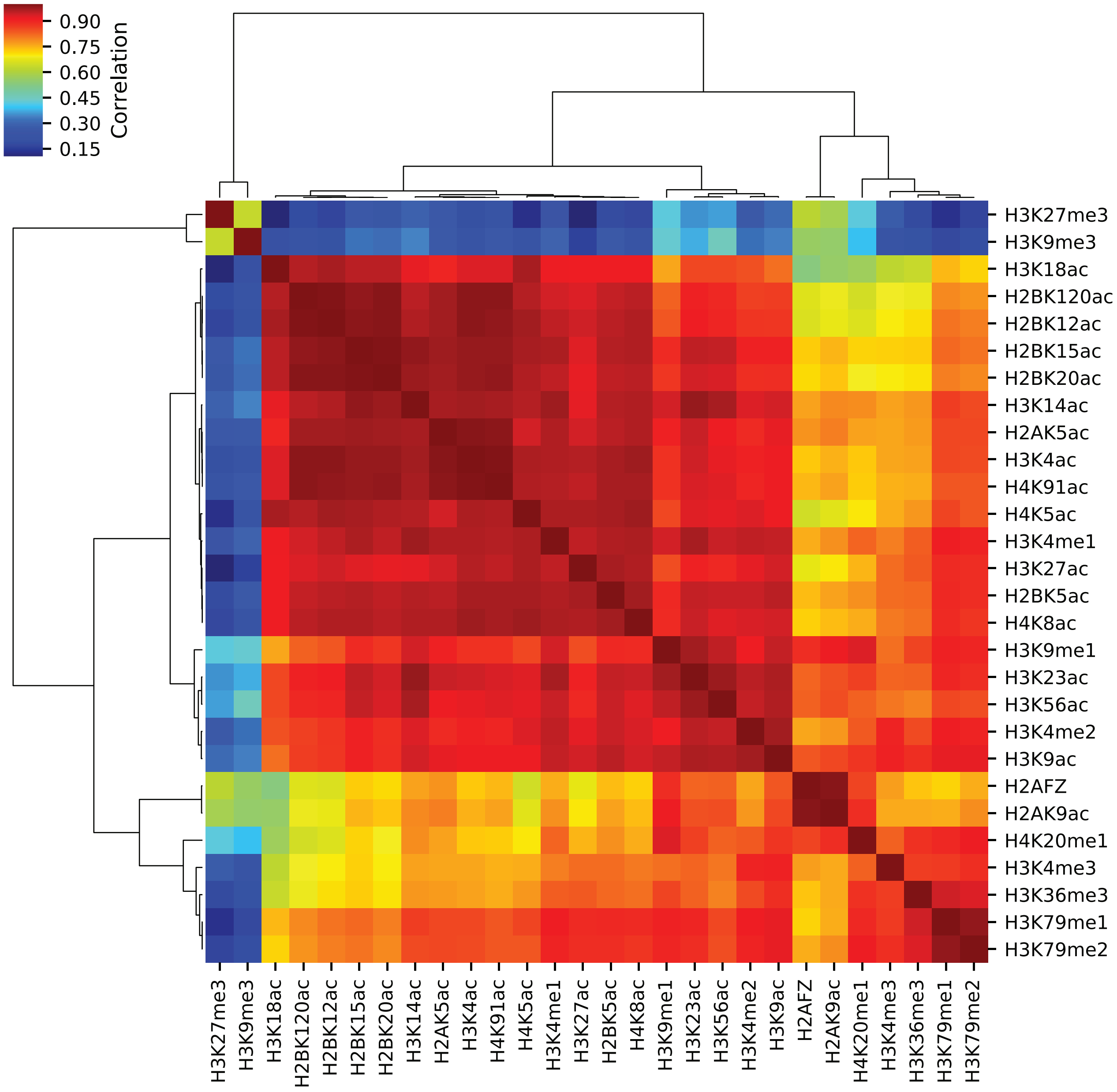


**Fig. S8.** Spearman correlation matrix of ChIP-seq signal distributions of the 28 histone post-translational modifications analyzed in this study across compartmental domains in IMR90 cells.

**Table S1. Values calculated from SAXS analysis of tandem bromodomains.**

| Protein | $R_g$ | $D_{max}$ |
|---|---|---|
| BrdT | 53.8 $\pm$ 1.2 | 188 $\pm$ 4 |
| Brd2 | 48.6 $\pm$ 2.9 | 181 $\pm$ 8 |
| Brd3 | 49.1 $\pm$ 2.6 | 183 $\pm$ 4 |
| Brd4 | 55.4 $\pm$ 5.5 | 200 $\pm$ 1 |
| Taf1 | 30.5 $\pm$ 1.2 | 103 $\pm$ 5 |

**Table S2. Brd4 tandem bromodomain ITC peptide binding parameters.**

| Peptide | $K_d$ | $N$ | $H$ (kcal/mol) |
|---|---|---|---|
| H4K5/8-diacetyl | 76.3 ± 6.4 | 1.09 ± 0.35 | -40.0 ± 1.8 |
| H4K5/8/12-triacetyl | 23.5 ± 1.2 | 1.22 ± 0.07 | -40.8 ± 2.9 |
| H4K5/8/16-triacetyl | 16.7 ± 1.7 | 0.73 ± 0.16 | -75.2 ± 17.7 |
| H4K5/8/20-triacetyl | 20.2 ± 0.5 | 0.92 ± 0.04 | -51.4 ± 2.4 |
